# More running causes more ocular dominance plasticity in mouse primary visual cortex: new gated running wheel setup allows to quantify individual running behaviour of group-housed mice

**DOI:** 10.1101/2025.04.25.647197

**Authors:** Cornelia Schöne, Jaya Sowkyadha Sathiyamani, Mihaela Guranda, Siegrid Löwel

## Abstract

Environmental enrichment boosts neuronal plasticity of standard-cage raised (SC) mice. Since it becomes increasingly more important to track individual mouse behaviours and its influence on brain plasticity, we designed a gated running wheel (gRW) setup allowing to correlate wheel running with neuronal plasticity, using the established paradigm of ocular dominance (OD)-plasticity after monocular deprivation (MD).

After SC-rearing until adulthood (>P110), group-housed mice were transferred to gRW cages, that provided an additional running wheel compartment for tracking individual wheel activity via implanted RFID chips. Notably, individual running parameters varied enormously: mice ran from close to 0 to ∼20 km across the 7 days of gRW experience, with on average running 0-3.96 km in 0-3.85 h/d and running bouts lasting from <1 up to 10 min, while running at a speed of 6-26 cm/s. OD-plasticity in V1 after 7 days of MD in the gRW was visualized using intrinsic signal optical imaging, and compared to control gRW-mice without MD via calculation of an OD-index. Most, notably - while wheel running enabled OD-plasticity -*individual* running parameters correlated with *individual* OD-indices after MD: Mice running longer distances, for longer time, at higher speeds and with longer and more frequent bouts displayed more experience-dependent V1-plasticity. In turn, a composite measure of overall running wheel activity derived from principal component analysis of running parameters accounted for 65% of inter-individual variability of OD-index following MD. Together our study demonstrates that interindividual variability of running behaviour is high, and mice intrinsically motivated to run more show enhanced V1-plasticity, underscoring the huge importance of analysing individual behavioural parameters together with any measure of brain plasticity. End

**Graphical abstract:** 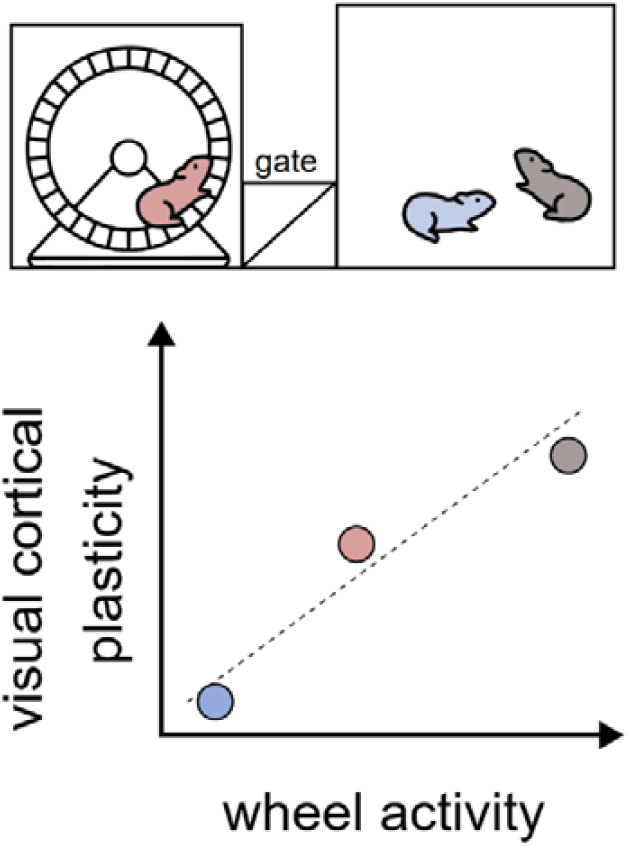

With our newly developed gated running wheel setup, we observed a striking correlation between individual running activity, and a measure of experience dependent plasticity in mouse primary visual cortex. More running caused more plasticity: running speed, running distance, total running time, number of running bouts and bout duration all correlated with a measure of visual cortical plasticity, the ocular dominance index. Thus, our observations add to the growing body of evidence that individual behavioural choices strongly affect individual brain plasticity.

## Introduction

While the natural environment of humans and mice is complex, that of laboratory rodents has been simplified and standardised in the past decades to an extent that deprives the animals of essential behaviours (Löwel et al., 2017; Cait et al., 2024). For instance, standard cage rearing (SC) leaves minimal room for the expression of individual traits and both behavioural and physiological interindividual variability is further reduced by using inbred strains lacking genetic variability (Wahlsten et al., 2006; Körholz et al., 2018). Recently, a number of laboratories demonstrated that raising animals under less deprived rearing conditions – in so-called “enriched environments”, i.e. housing in larger social groups and/or larger cages, with regularly changed mazes to navigate through and running wheels for voluntary physical exercise – elicits remarkable effects on brain wiring and plasticity across molecular, anatomical, and functional levels when compared to animals raised in an SC environment (Sale et al., 2007; Fabel et al., 2009; Baroncelli et al., 2010; Di Garbo et al., 2011; Greifzu et al., 2014; Kalogeraki et al., 2014; Löwel et al., 2017; Stryker and Löwel, 2018; Bogado Lopes et al., 2023).

Both environmental enrichment, but also just voluntary wheel running have been shown to boost experience-dependent changes in rodent primary visual cortex (V1) (Sale et al., 2007; Baroncelli et al., 2010; Greifzu et al., 2014; Kalogeraki et al., 2014; Kaneko and Stryker, 2014), using the established model of ocular dominance (OD) plasticity after monocular deprivation (MD)(Gordon and Stryker, 1996; Cang et al., 2005): when one eye is closed for few days, the relative V1-activation strength is shifted towards the open eye (Cang et al., 2005; Gordon and Stryker, 1996). SC-mice beyond P110 do no longer show OD-plasticity with 7 days of MD (Lehmann and Löwel, 2008; Sato and Stryker, 2008), and need ∼7 weeks for V1-activation changes (Hosang et al., 2018)

While it is clearly established that running increases OD-plasticity in mice (Kalogeraki et al., 2014; Kaneko and Stryker, 2014) the consequences of interindividual variability of wheel running on experience-dependent V1-plasticity has not yet been studied. We therefore aimed to analyse the effects of individual running behaviour on OD-plasticity of adult (>P110) group housed SC-mice. As tracking individual wheel running activity of group housed mice is still challenging (Mayr et al., 2020; Reuser et al., 2022), we designed a custom-built gated running wheel (gRW) setup consisting of a rat-sized home cage connected to a compartment with a surveyed running wheel that allowed individual tracking of mice implanted with an RFID sensor. We quantified running distance, time, speed, bout number and bout length of running activity and tested whether they correlated with the OD-index, which compares visual stimulus evoked V1-activation via the ipsi- and contralateral eye and thus quantifies the magnitude of OD-plasticity (Cang et al., 2005).

Running parameters varied extensively between individual mice, with some mice running only few meters while others ran >20 km during their 7 days of running wheel exposure. Most notably, correlating the macro (running distance, time and speed) and micro (bout number and duration) architecture of running wheel activity of individual mice with their individual OD-index, quantifying OD-plasticity, revealed strong correlations with Pearson correlation coefficients ranging from -0.77 to -0.83: more active runners displayed stronger OD-plasticity. Our results underscore the importance of taking individual behavioural choices into account when analysing influences on brain plasticity.

## Methods

### Animals and rearing conditions

In order to avoid conflict between group housed males, we limited our experiments to groups of 2-5 adult female C57Bl6/J mice (age range P127-225, called P160), growing up in Tecniplast 1284 standard mouse cages (SC; size: 17x32x19cm) until the day an MD/noMD was performed. After the MD/noMD, animals were cohoused in the gRW-setup, provided with *ad libitum* food and water, and maintained on a 12 h light dark cycle.. All experimental procedures complied with the National Institutes of Health guidelines for the use of laboratory animals and were approved by the local government of Lower Saxony, Germany (Niedersächsisches Landesamt für Verbraucherschutz und Lebensmittelsicherheit).

### Optomotry, monocular deprivation and implantation of RFID tags

To ensure that all experimental mice had regular vision before testing OD-plasticity with MD, we measured the spatial frequency threshold (SFT) of the optomotor reflex of both eyes using an optomotor system (Prusky et al. 2004; Lehmann and Löwel, 2008). SFT values were within published values and differed by less than 0.02 cyc/deg between the two eyes in all mice.

Thereafter, the right eye was sutured shut (MD) for 7 d according to published protocols (Gordon and Stryker, 1996; Cang et al, 2005; Greifzu et al., 2014). In brief, mice were anaesthetized using 2.5 % isoflurane in 1:1 O_2_/N_2_, and kept at 1-1.5% isoflurane for stable anaesthesia. Body temperature was maintained at 37°C using a heat pad and rectal probe. After subcutaneous injection of Rimadyl (Carprofen, 5 mg/kg) and covering of eyes with Bepanthen eye cream, a local analgesic (Lidocaine, xylocaine gel 2 %, Aspen Pharma Trading Limited, Ireland) was applied to the eyelid of the right eye before trimming and closing of the eye with 2 mattress stiches (suture material: 7-0 Perma-Hand silk, Ethicon, 8.0mm diameter). The eyes of the animals were examined daily to ensure that the MD-eye stayed closed. Additionally, optomotor reflex thresholds (SFT) of the open eye were tested daily after MD, as a measure for functional MD, including the day of intrinsic signal optical imaging.

Cylindrical glass-covered radio frequency identification devices (RFID; length: 12.5 mm; diameter: 1.93 mm, Sparkfun Electronics, Colorado, USA) were implanted during the MD or noMD (control without MD) surgery to minimize mouse discomfort. First, a patch of fur was shaved in the animal’s neck using small surgical scissors, then a local analgesic gel (Lidocaine, xylocaine gel 2 %, Aspen Pharma Trading Limited, Ireland) was applied to the exposed skin. A small incision was made and the chip carefully placed subcutaneously in the scruff to make it difficult for the mice to pick at and thereby reducing the risk of an infection. The incision was sealed with a simple interrupted stitch (suture material: 7-0 Perma-Hand silk, Ethicon, 8.0mm diameter). After MD/RFID implantation and complete recovery from anaesthesia, mice were moved to a gRW setup.

### Gated running wheel setup (gRW)

In order to assess individual running wheel behaviour, we developed an open source gated running wheel (gRW) cage, which served as the home cage for the duration of the experiment (Figure 1A, Table 1). It consisted of a standard size rat cage (43cm x 27cm) connected to a separate gRW-compartment, equipped with the same RW as used in the enrichment cages in Greifzu et al., 2014 (Figure 1A). Wheel diameter was 12.5 cm, so that the calculated RW-circumference was 39.3 cm. RW-turns were registered using a hall sensor for detecting motion of two magnets attached at opposing sites of the RW. The mice needed to pass through a seesaw in order to reach the RW-compartment. Entering would flip the seesaw blocking other mice from entering (see video 1 and 2). To ensure the seesaw remained closed while a mouse was inside the RW-compartment, the seesaw was actively blocked from flipping back. When activity was detected by beam break sensors outside the RW-compartment, an electromagnet was activated to hold the seesaw closed. Attempts of mice to exit the RW-compartment were detected using beam break sensors inside the RW-compartment, which would deactivate the seesaw magnet. Seesaw position was registered through a roller switch.

**Table 1:** Hardware essentials for building a gated running wheel setup.

| Item | Catalog No. | Company |
| --- | --- | --- |
| Raspberry pi 4B+ 4GB, 32 GB SD | RP-4B-4GB | Raspberry Pi Foundation, Cambridge, England |
| Raspberry Pi Noir Camera 2.0 8MP | RB-CAMERAV2IR | Raspberry Pi Foundation, Cambridge, England |
| Raspberry Pi® RB-LCD-7 Display-Modul 17.8 cm (7 Zoll) 800 x 480 Pixel | RB-LCD-7 | Raspberry Pi Foundation, Cambridge, England |
| RFID reader | ID-20LA | SparkFun Electronics, Colorado, USA |
| RFID implants | SEN-09416 | Sparkfun Electronics, Colorado, USA |
| roller switch for seesaw | 190.072.013 | Marquardt, Germany |
| phototransistor for beam break sensors | BPV11F | Vishay, Germany |
| reed contact for Hall sensor | MS-213-3 | PIC Proximity Instrument Controls GmbH, Germany |

**Figure 1.**
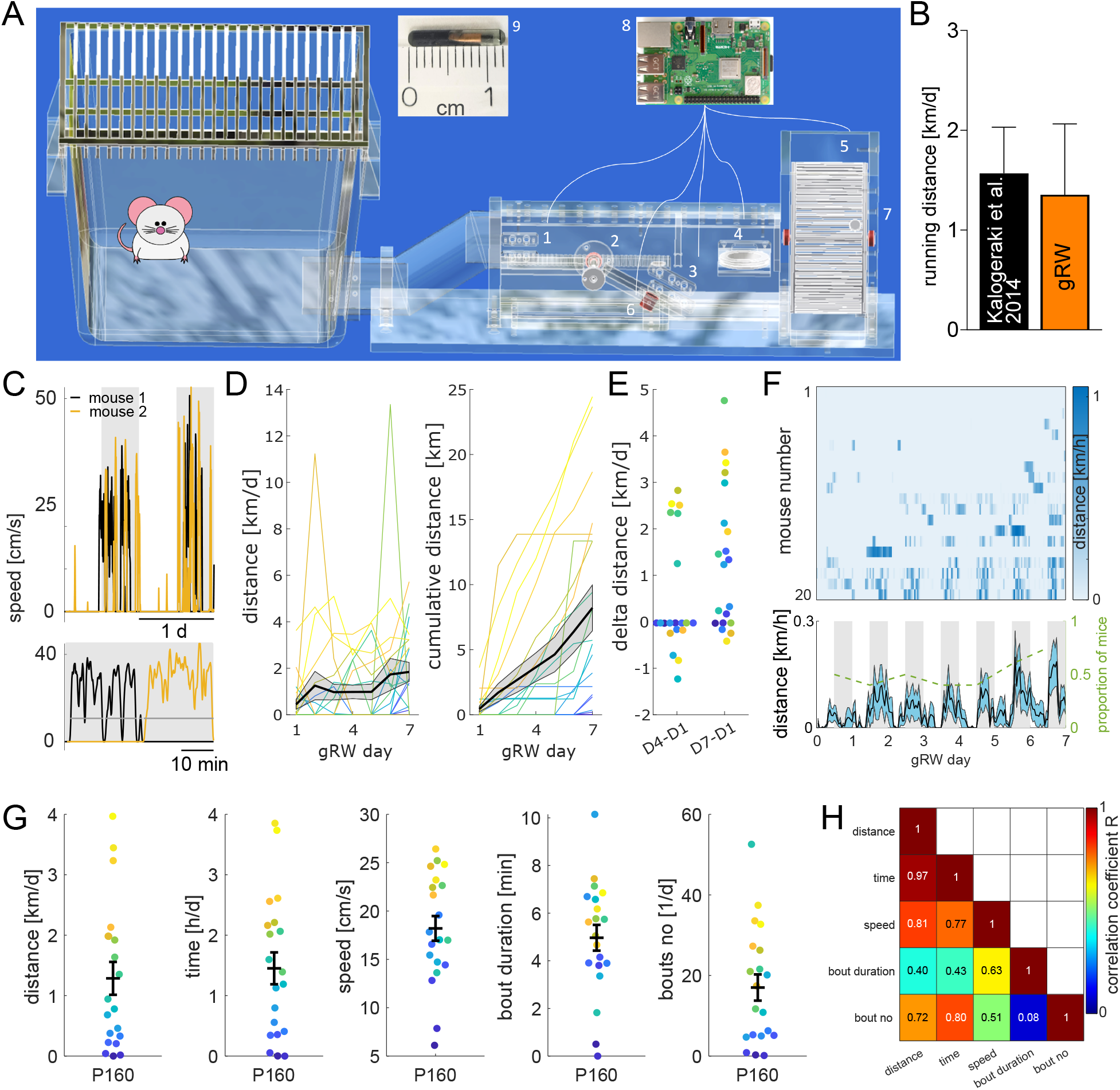
Gated running wheel setup allows to track individual mouse running wheel activity. **A**. Standard-sized rat cage (left, 43 cm x 27 cm) with an additional compartment (right) containing beam break sensors (1,3), a seesaw (2), RFID sensor (4), hall sensor (5), magnet (6), and a running wheel (7). All the sensors are connected to a raspberry pi 4B+ (8). (9) Illustrates the size of the implanted RFID. **B**. Comparison of average daily running activity of gRW housed mice (orange) with published data from Kalogeraki et al. 2014 (black). **C**. Example of quantified gRW-activity of two co-housed mice illustrating allocation of running bouts to individual mice (orange/black line) for 2 consecutive days and nights (grey background) (top) and 60 minutes during night time (bottom), with detected running bouts highlighted in grey. **D**. Daily (left) and cumulative (right) running trajectories of individual mice housed in gRW enrichment for 7 days, document a large range of individual running behaviour. While some mice achieve high distances from the first day, for most mice daily running distances only achieved high distances torwards the end of the 7 days. Colours of traces and dots are matched for individual mice across figures E-G **E**. Overall there was a significant increase in daily running between day 1 and day 7, but not day 1 and 4. **F**. Running distances of all animals across 7 days binned per hour with top: heat map of indivdual mice and bottom: average distances across all animals, with overlaid proportion of mice running on each day (defined as at least 1 bin above 0 distance travelled). Note that animals with sufficient running activity predominantly run during night time (higher average values and darker patches in heatmap), with few expceptions of running also during the light period. **G**. Running wheel parameters in individual mice. Note the huge interindividual variability across parameters. Mice that ran longer distances also spent more time running at higher speeds with larger bout numbers. **H**. Correlation matrix illustrating interdependencies of 5 running wheel parameters displayed in G.

Mice were chipped with radio frequency identification devices (RFID). The RFID-tags transmitted the encrypted individual ID to a USB-driven reading device (ID-20LA SparkFun Electronics, Colorado, USA) at 125 kHz via a custom-made RFID coil. Thus, each time a mouse passed underneath the RFID coil into the gRW compartment, RFID and time stamps were saved. All sensors were connected to the GPIO pins of a raspberry pi 4B+. In addition, a raspberry pi noir camera was used to record movement inside the RW-compartment at 40 Hz; camera data were used to confirm sensor data and to extract RW-turns when the Hall sensor failed. All data was recorded using custom written python scripts running on the raspberry pi. A 3D .pdf file illustrating dimensions of cage components and electronic assembly is provided under supplementary material (Model 1 and Figure S1, respectively).

### Analysis of gated wheel running

In order to assign each RW-turn to individual mice, we first defined running bouts as running at a speed above 0.1 Hz, then the most recently detected RFID was assigned to all wheel turns within the respective running bout. When more than one RFID was detected since the last gate flip, this indicated that more than one mouse had entered the gRW compartment. This occasionally happened, due to two mice squeezing through the gate simultaneously, or due to failure of the magnetic seesaw lock. In this case, the RW-turns were labelled as ambiguous. Mice with ambiguous RW-turns of more than 7% were excluded from the RW-analysis, but included for analysis of OD-plasticity in Figure 2. For the remaining mice, on average, we quantified 1.7±0.5% ambiguous RW-turns (n=20 mice), confirming that the self-locking seesaw of the gRW was well suited to restrict access to the wheel compartment to individual mice.

**Figure 2.**
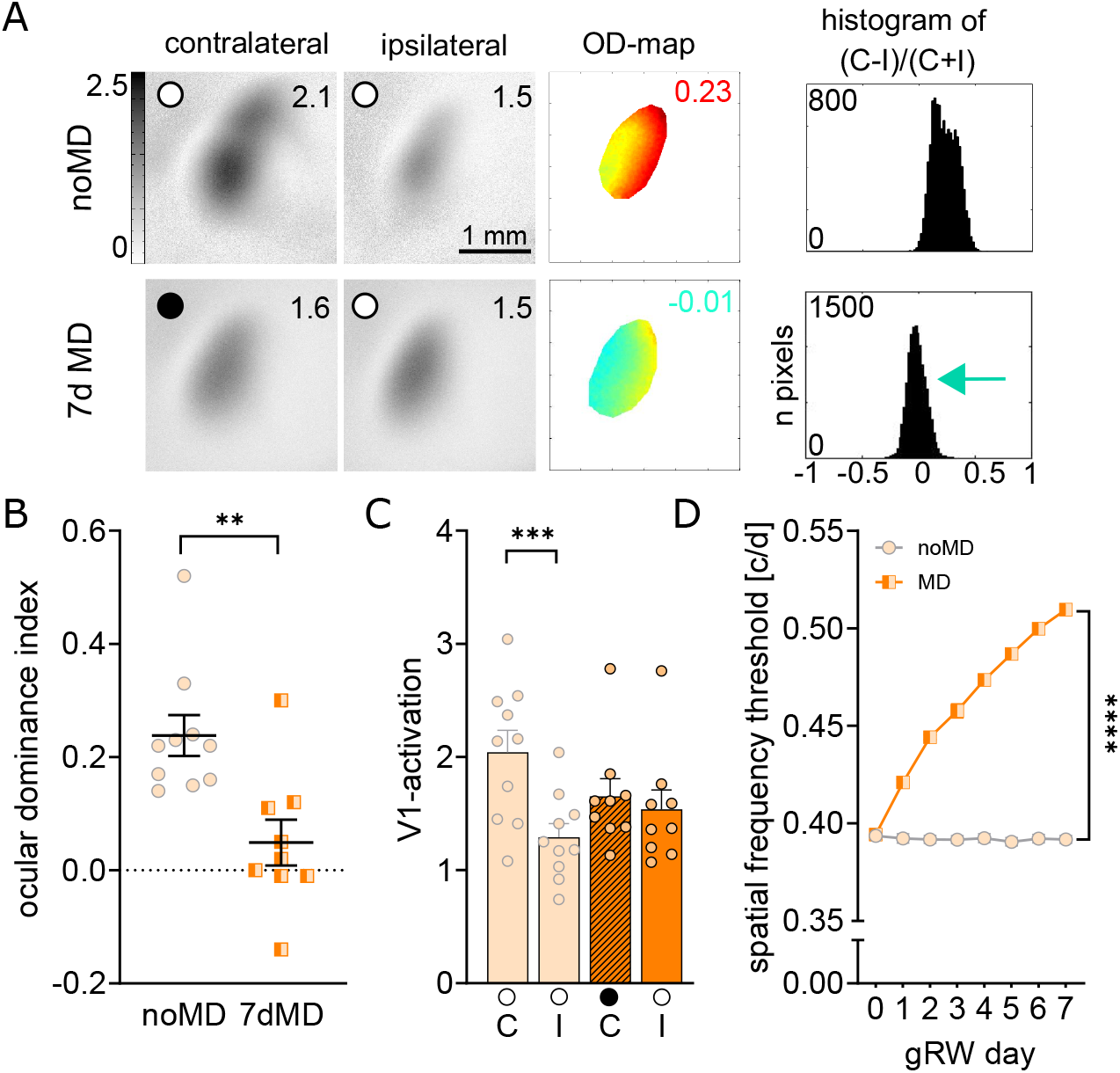
gRW enrichment boosts ocular dominance plasticity in group-housed mice. **A**. Intrinsic signal optical imaging examples of grey-scale coded activity maps after visual stimulation of the contralateral and ipsilateral eye in binocular V1 of C57Bl/6J gRW housed mice after 7 days of monocular deprivation (MD) and without MD (no MD). The magnitude of V1-activation is expressed as the fractional change in reflectance x10^-4^ and indicated in the top right. Warm colors in the color-coded OD-maps indicate contralateral eye dominance, colder colors indicate an OD-shift towards to ipsilateral eye. Histograms in A,E represent distribution of pixel wise OD-scores. Scale bar: 1mm. **B**. Quantification of OD-index. Symbols represent values of individual mice, means are marked by horizontal lines. **C**. V1-activation elicited by stimulation of the contralateral (C) or ipsilateral (I) eye (filled circle indicates MD eye). Note that MD did not reduce V1-activation through the deprived eye in C. **D**. Spatial frequency threshold of the optomotor reflex for noMD and MD mice, plotted vs. days (x-axis). Note that thresholds increase after MD, indicating successfully induced MD (*p < 0.05, **p < 0.01, ***p < 0.001 and ****p < 0.0001 in Mann-Whitney or 2-way ANOVA with Sidak’s post hoc test).

### Optical imaging of intrinsic signals and visual stimuli

As a final step of the experiment, OD-plasticity was assessed after 7 days of MD/noMD in the gRW-setup, using optical imaging of intrinsic signals (Cang et al., 2005; Greifzu et al., 2014).

#### Surgery

Briefly, mice were box-anaesthetised with 2.5% halothane in O_2_ and N_2_O (1:1) and injected with atropine (5mg/kg, s.c.; Franz Köhler Chemie), dexamethasone (8mg/kg s.c.; Merck), and chlorprothixene (8mg/kg, i.m.; Sigma-Aldrich). After placing animals in a stereotaxic frame, anaesthesia was maintained with 0.8% halothane in a 1:1 mixture of O_2_ and N_2_O.

#### Data acquisition and visual stimulation

(Kalatsky and Stryker, 2003; Cang et al, 2005). Mouse V1 responses were recorded through the skull using the “Fourier” imaging method of Kalatsky and Stryker (2003) and optimised for the assessment of OD-plasticity (Cang et al., 2005). V1-signals were visualised with a CCD camera (Dalsa 1M30) using a 135x50 mm tandem lens configuration (Nikon) with red illumination light (610±10 nm). Active brain regions absorb more red light and appear darker in the images. Frames were acquired at a rate of 30 Hz, temporally binned to 7.5 Hz, and stored as 512x512-pixel images after spatial binning of the camera image. Visual stimuli were presented on a 60 Hz refresh rate monitor (21 inches; Accuvue HM-4921-D, Hitachi) positioned 25 cm from the eyes. Stimuli consisted of white drifting horizontal bars (2° wide) limited to the contralateral binocular visual field (-5° to 15°) as described previously (Greifzu et al., 2014). The amplitude component of the optical signal represents the intensity of neuronal activation (expressed as fractional change in reflectance times 10^-4^) and was used to calculate an ocular dominance (OD) index. At least three maps per animal were averaged by an experimenter blinded to the experimental conditions to compute the OD-index as (C+I)/(C-I), with C and I representing the response magnitudes of each pixel to visual stimulation of the contralateral (C) and ipsilateral (I) eye.

## EXPERIMENTAL DESIGN AND STATISTICAL ANALYSES

### Principal component analysis

Principal component analysis was performed using the MATLAB (R2022b) pca function on wheel running parameters (distance, time, speed, bout duration and bout number) obtained from combined noMD and MD mice in order to eliminate multicollinear variables which effectively reduces dimensions of descriptive variables with the first principal component (PC1) capturing the maximum variance of the data (86-90% of variance in wheel running parameters, depending on whether average or individual data across days was used, respectively). In both cases, PC1 had the biggest loading for bout no., with smaller loadings for all other wheel parameters (Figure S2). Together this suggests, that PC1 is well suited to represent overall wheel activity capturing maximal variance of individual running wheel performance.

### Statistical analysis

Normality of data was assessed using the Kolmogorov-Smirnov test, and non-parametric versus parametric tests were chosen accordingly as stated in the text. Inter-group comparisons between two groups were done by Student’s t-test or Mann Whitney test, for 3 or more groups, we used one or two-way analysis of variance (ANOVA), as detailed. In analyses in which a within-subject factor was present (i.e. eye), ANOVA with repeated measurements was performed. Post hoc multiple comparison tests were corrected by applying Sidak correction to p-values. The levels of significance were set as *: p<0.05; **: p<0.01; ***: p<0.001. Data are represented as means ± standard error of mean (s.e.m.).

### Software accessibility

Any software and code used for running the gRW setup and for processing obtained data are available upon request.

## Results

### The gated running wheel setup allows for detailed readout of individual mouse running behaviour

Given the importance of social interactions for the well-being of mice, we specifically designed a gated running wheel setup (gRW), so that we could track individual running activity of group-housed mice in a home cage setting. The gRW setup consists of a standard rat cage connected to a separate compartment with a running wheel for voluntary physical exercise (Figure 1A). Upon entering the gRW-compartment, mice are individually registered using an implanted RFID sensor allowing to allocate running bouts to individual animals. This enables us to quantify individual wheel running parameters and to correlate individual running behaviours with individual V1-plasticity parameters assessed using intrinsic signal optical imaging.

In order to provide a detailed analysis of wheel activity of mice, we first compared running wheel activity between animals undergoing monocular deprivation (MD, n=12) and control mice (noMD, n=8): wheel running was not affected by MD (Figure S2). Next, we evaluated whether the separate gRW-compartment would impair access and reduce overall running activity compared to a previous publication using identical running wheels located inside the home cage. This did not seem to be the case: mice in the gRW setup ran 3307±763 RW turns per day, corresponding to 1.3±0.3 km/d (Figure 1B), rather similar to the 3991±445 RW turns/day, corresponding to 1.6±0.2 km/d of the animals of a previous study (Kalogeraki et al., 2014).

Since, typically, only a single mouse entered the RW compartment at a time, the activity of individual mice could be well separated by assigning running bouts to individual mice for assessing interindividual variability in wheel running (Figure 1C-G). Across mice, running distance increased over days as revealed by linear regression analysis (Figure 1D, F(1,138)=4.98, p=0.027). However, running was not consistent across days for individual mice (Figure 1D), and could range from close to 0 to above 10 km on individual days and for individual mice (Figure 1D). Average cumulative running distances reached 8.2±1.8 km after 7 days, ranging from close to 0 to above 20 km of total distance travelled by individuals (Figure 1D). Mice exhibited a significant increase in running distance from day 1 to day 7 (Figure 1E, one-sample t-test p=0.001), whereas the difference between day 4 and day 1 was not significant (Figure 1E). This was paralleled by a larger proportion of mice running more than one bout per day, which increased from 50% on day 1 to 75% on day 7 (Figure 1F).

As expected from nocturnal animals, mice ran more during night time compared to day time, with approximately 50% of mice showing a clear circadian pattern and running predominantly during night time (Figure 1F), while the other half of the animals did not run sufficiently to detect circadian activity preferences, and few individuals also occasionally ran during day time.

On average, mice achieved running times of 1.5±0.3 h/d consisting of 17±3 bouts/d with individual running bouts lasting on average 5.0±0.5 min, while running at a speed of 18±1 cm/s (Figure 1G). Of note, mice covered a wide range of running wheel parameters, running 0-3.96 km/d in 0-3.85 hours with running bouts lasting from <1 up to 10 min, while running at a speed of 6-26 cm/s. Interestingly, mice running longer distances typically also spent more time running, at faster speeds, which was also reflected in a larger number of running bouts, with few exceptions, as illustrated by the colour coding of individuals in figures 1E,G. A correlation matrix was computed to examine the interrelationships between the 5 quantified running parameters more precisely (Figure H): The strongest correlation was observed between distance and time (R=0.97, p<0.0001), i.e. mice that ran for longer time periods also travelled a larger total distance. Speed and bout number also strongly correlated with running distance (speed: R=0.81, p<0.0001; bout no: R=0.72, p<0.0001), i.e. mice running faster and more often, also reached a higher final distance. In turn, bout duration did not correlate with distance, time or number of bouts (p>0.05), however there was a significant correlation with running speed (R=0.63, p<0.01), i.e. faster mice also ran longer bouts.

Together, these data demonstrate i) substantial interindividual variability in multiple running wheel parameters of our group-housed mice which may influence cortical plasticity, and ii) that our new gRW-setup is ideally suited to analyse this question in detail.

### Gated wheel running restores OD-plasticity to adult standard cage raised mice

We had previously shown that running can boost OD-plasticity in SC-raised adult mice, even if running was possible only during 7 days of MD (Kalogeraki et al., 2014). Here, we tested whether i) the gRW setup also boosts OD-plasticity in SC-mice and ii) individual running parameters, i.e. running more, would influence individual OD-plasticity. gRW-housing started immediately after MD, and V1-activity maps were visualized after 7 days of MD. We recorded intrinsic signal optical imaging responses of binocular V1 to visual stimulation of the left and right eye with horizontal drifting bars (Cang et al., 2005).

Confirming previous imaging data in mice without MD (Lehmann and Löwel, 2008; Sato and Stryker, 2008), visual stimulation of the contralateral eye induces a stronger V1-activation compared to ipsilateral eye stimulation, visible as darker patches in the V1-activity maps (Figure 2A), and reflected in a higher V1-activation strength (Figure 2C; contra/ipsi: 2.0±0.2/1.3±0.1, n=10, p=0.0003). In contrast, in gRW mice, MD induced a clear OD-shift: After MD, the contralateral eye no longer activated V1 more strongly than the ipsilateral eye (contra/ipsi: 1.7±0.2/1.5±0.0.2, n=9; p=0.7). Calculating the OD-index, quantifying OD-plasticity by comparing ipsi- and contralateral eye induced V1-activity (Fig. 2B), confirms the gRW boosted OD-shift: in noMD mice, the OD-index was 0.24±0.04 (n=10), indicating contralateral eye dominance, whereas following MD, the OD-index was reduced to 0.05±0.04 (n=9, Mann-Whitney p-value 0.001), demonstrating a clear OD-shift in P160 mice. Thus, our new gRW setup boosts OD-plasticity in group-housed mice.

### Optomotor results

To test animals’ basic visual abilities and to confirm the effectiveness of MD, we used the virtual reality optomotor system, and measured the spatial frequency threshold (SFT) of the optomotor reflex before and after MD (Prusky et al., 2004): vertical drifting gratings of various spatial frequencies evoke small horizontal head and neck movements following the stimulus. MD typically leads to an enhancement of optomotor reflex thresholds through the open eye (citation!). Confirming effective MD, the SFT of our group-housed mice increased from 0.39±0.00 cyc/deg on day 0 to 0.51±0.00 cyc/deg on day 7 after MD (n=10/9; 2-way ANOVA for effect of MD p< 0.0001; Figure 2H), while values remained stable in noMD mice (day 0/7: 0.39±0.00/0.39±0.00 cyc/deg, 2-way ANOVA for effect of MD: p<0.0001).

### Gated running wheel parameters correlate with a measure of ocular dominance (OD) plasticity, the OD-index

#### Correlation of overall running wheel activity with OD-index

In order to investigate how individual behavioural choices shape brain plasticity, we tested whether individual running wheel parameters were correlated with individual OD-indices, which quantify the magnitude of OD-plasticity. For this, we first obtained a measure of overall gRW activity for individual mice by performing a principal component analysis (PCA) on the quantified running wheel parameters (distance, time, speed, running bout numbers, bout duration) of pooled no MD and MD mice, averaged across the 7 days of wheel exposure. The first principal component explained 87% of interindividual variability in wheel running and hence was a suitable measure of overall gRW activity of individual mice (see methods and figure S2 for details). Strikingly, interindividual variability in gRW activity explained 65% of variability of the OD-index (r^2^=0.65, p=0.017, Figure 3A,B): Animals with high gRW performance showed a lower OD-index and thus increased plasticity (Figure 3A,B). In contrast, correlation analyses in no MD mice revealed no significant relationship between gRW activity and OD-index (r^2^=0.02, p=0.77, Figure 3B).

**Figure 3.**
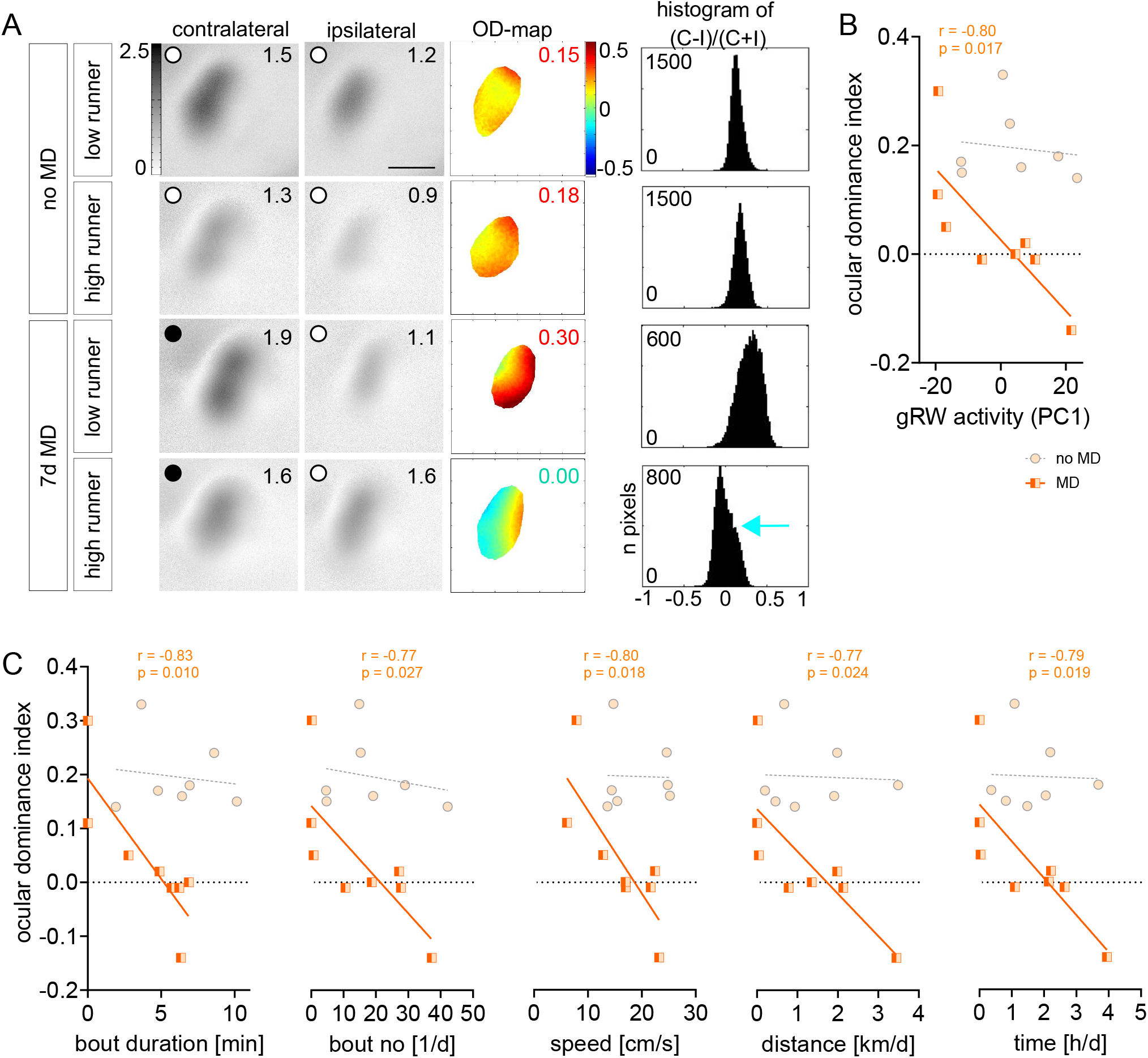
Individual running wheel (RW) activity correlates with ocular dominance (OD) plasticity, quantified via the OD-index after monocular deprivation (MD) in P160 mice. **A**. Example intrinsic signal optical imaging recordings from noMD and MD P160 animals, with low (gRW activity <0) and high (gRW activity >0) RW performance (gRW activity was defined as the first principal component (PC1) of RW parameters displayed in C, and shown in B). Data displayed as in figure 2A. Note that OD-plasticity after MD is only boosted in P160 mice, when mice run sufficiently in the gRW setup. **B**. The OD-index of individual P160 mice is correlated with their individual gRW activity after MD (orange), but not in no MD control mice (grey). Lines represent linear regression fitted to MD mice and no MD mice, respectively. Pearson r-values and p-values for MD mice are in the figure. **C**. The OD-index of individual mice after MD is correlated with all quantified running wheel parameters. Data is displayed as in B. Pearson correlation between OD-index and individual wheel running activity was significant for all tested running wheel parameters in P160 mice after MD, but not no MD control mice.

#### Correlation of individual running wheel parameters with OD-index

In addition, we also observed striking correlations of individual running wheel parameters with individual OD-index after MD (Figure 3C). In MD mice, bout duration, speed, bout number, running distance and running time explained close to or above 60% (e.g. r^2^ ≥ 0.60) of the interindividual variability in OD-index of MD mice (n=9; bout duration: r^2^=0.69, p=0.0102; speed: r^2^=0.63, p=0.0175; bout no: r^2^=0.58, p=0.0271; running distance: r^2^=0.60, p=0.0243; running time r^2^=0.62, p=0.0192). Importantly, no significant correlation between running wheel parameters and OD-index was found in the no MD control group (p>0.05 for all parameters), suggesting that baseline OD-index is not affected by wheel running.

#### Correlation of wheel activity on specific days with OD-index

Previous experimental evidence has demonstrated that running wheel activity correlates with cortical gamma activity in V1 (Niell & Stryker, 2010), and that MD is associated with increased gamma activity in V1 for several hours after MD, at least in juvenile mice aged 24-27 days (Quast et al., 2023). Hence, we wondered whether running wheel activity on a specific day after MD was correlated more strongly with the OD-index. For this we used PCA for dimensionality reduction of running wheel parameters measured for *each day separately* and correlated the first principal component (explaining 90% of variability of running wheel parameters across days and animals) with the OD-index: gRW activity significantly correlated with the OD-index on days 6 and 7 after MD (day6/7: r^2^=0.52/0.53, p=0.045/0.041), while this relationship was not significant on days 1-5 (p>0.05 for all correlations), suggesting that predominantly running on days 6 and 7 affects the OD-index. Nevertheless, gRW activity on the first days still contributed to individual variability in OD-index, since including gRW activity of cumulative numbers of days in the analysis already produced significant correlations when only data from days 1 and 2 was included (p>0.05 for day 1 alone, p<0.05 for data from >1 days).

Overall these data document that group housed mice display huge interindividual variability in voluntary physical exercise when given the possibility to use a running wheel: individuals differ in bout number, bout duration, running speed, running distance and running time. Most notably, the amount of wheel running was predictive of the magnitude of OD-plasticity: Mice that ran more had stronger experience-dependent changes in V1-activation after MD. This directly highlights the importance of inter-individual behavioural variability for brain plasticity and suggests that individual behavioural choices and their influence on brain physiology and plasticity should be an integral part of future studies.

## Discussion

Using our newly developed gated running wheel setup (gRW) that allows to track individual running parameters of group-housed mice, we observed an enormous variability of individual behaviours of the tested animals: running speed, running distance, total running time, number of running bouts and bout duration varied by many orders of magnitude between individuals. Furthermore, we revealed a striking correlation between individual running parameters and experience-dependent plasticity in mouse V1. More running caused more ocular dominance (OD) plasticity: all quantified running parameters significantly correlated with a measure of visual cortical plasticity, the OD-index. Thus, our observations add to the growing body of evidence that individual behavioural choices can strongly affect individual brain plasticity, and should therefore be considered when analysing neuronal plasticity.

Mice housed in the gRW for one week showed huge interdividual differences in running wheel performance, running 0-3.96 km/d in 0-3.85 hours with running bouts lasting between 0 and 10 min, while running at a speed of 6-26 cm/s, suggesting diverse intrinsic motivation to run. Wheel running is distinct from home cage running because the distances and speeds reached per day are drastically increased when mice have access to a running wheel. Mice in their home cages only achieve speeds of up to 1 cm/s and distances of 0.1-0.2 km/d (Iannello, 2019), while for running wheels allow speeds of up to 15-120 cm/s and distances of 1-20 km/d have been reported (Koteja et al., 1999; Kopp, 2001; Manzanares et al., 2018). While our data spans the lower end of the reported range - with individual mice running up to 10 km/d on single days – this is expected from the specific conditions used here: i) group housing reduces running distances by ∼50% (Plenz and Kanold, 2021) and ii) the short-term wheel access provided here for only 7 days likely was not sufficient for mice to reach a stable performance, which has been reported to take ∼2 weeks (De Bono et al., 2006).

It is widely accepted that OD-plasticity in SC-raised mice is age-dependent, with clearly decreasing plasticity in adult animals beyond P110, which requires an extended MD for observable OD-shifts (Sawtell et al., 2003; Pham, 2004; Hofer et al., 2006; Lehmann and Löwel, 2008; Sato and Stryker, 2008; Hosang et al., 2018). A number of environmental and behavioural interventions have been established by now for restoring plasticity in adult SC-raised rodents or for sustaining the plastic potential of adult rodent V1 beyond that age (Espinosa and Stryker, 2012; Hübener and Bonhoeffer, 2014). This includes previous episodes of MD (Hofer et al., 2006), forced visual stimulation (Matthies et al., 2013), combined visual stimulation and running (Kaneko and Stryker, 2014), and dark rearing (He et al., 2006; Stodieck et al., 2014), all of which promoted visual cortical plasticity in adult animals. Notably, mice and rats with access to a running wheel or housing in enriched environment retained OD-plasticity into late adulthood (Sale et al., 2007; Baroncelli et al., 2010; Greifzu et al., 2014; Kalogeraki et al., 2014), demonstrating that voluntary physical exercise can sustain the brain’s capacity for adaptive modification to environmental changes.

While OD-shifts obtained in the present gRW-mice were similar to previously published data after 7 days of RW-enrichment (Kalogeraki et al., 2014), OD-shifts were smaller compared to animals experiencing lifelong enrichment in even larger 2-floor cages with a regularly changed maze (Greifzu et al., 2014). Thus, while short-term running can restore OD-plasticity in adult mice, more complex enrichment of their immediate environment for longer periods of time, enables stronger experience-dependent V1-activity changes. Alternatively, extension of MD-duration can also boost OD-plasticity in adult SC mice (Hosang et al., 2018). Thus, there seems to be a trade-off between age and MD duration in SC-raised animals: in younger mice, shorter MDs are sufficient to induce significant OD shifts. In contrast, older SC-mice need considerably extended MD duration to display OD plasticity, but these long MD times can be shortened by specific environmental and behavioural interventions such as, e.g., raising animals in an enriched environment (Sale et al., 2007; Greifzu et al., 2014) or providing access to a running wheel (Kalogeraki et al., 2014), like in the present study.

Using C57Bl/6J inbred mice housed in the gRW-setup allowed us to test how individual behavioural trajectories affect brain plasticity with minimal influence of genetics on the observed phenotype. In line with the model of Kempermann (2019), we observed a linear relationship between running parameters and OD-plasticity quantified by the OD-index, with individual running performance predicting up to 65% of phenotypic variability in OD-index between mice. Thus, individual behavioural choices exert a strong influence on experience-dependent plasticity. Interestingly, all wheel parameters correlated similarly strong with individual OD-index, with p-values of correlations only varying slightly between 0.010 and 0.027 between bout duration, bout number, running speed, running distance and running time. As wheel running comprises a rewarding behaviour that even mice in the wild pursue voluntarily (Sherwin, 1998), it remains unclear why some mice showed very little wheel activity. Multiple factors affecting wheel running have been identified, including mouse strain, sex, group housing, social hierarchy, age and circadian rhythms (Kopp, 2001; De Bono et al., 2006; Basterfield et al., 2009; Bartling et al., 2017; Bains et al., 2018; Balog et al., 2019; Plenz and Kanold, 2021). Importantly, social hierarchy has been shown to affect both wheel running and OD-plasticity in male mice, with dominant males exhibiting stronger OD-shifts compared to subordinate males (Balog et al., 2019). While social hierarchies are less prominent in female mice (Williamson et al., 2019) used here, we cannot exclude that subordinate social rank may have caused reduced access to the gRW in some of our mice, and thus might have contributed to both the amount of wheel running and variability of OD-index.

Both high running speed and bout duration require high physical fitness and might be associated with better sensory-motor coordination for efficient running that may improve with practice (De Bono et al., 2006), suggesting that pre-training or rearing under less deprived conditions could also benefit OD-plasticity by enabling more efficient running and higher fitness. Interestingly, as intermitted access to running wheels has been shown to specifically increase adult neurogenesis in the hippocampus as opposed to continuous long term access (Nguemeni et al., 2018), gRW housing might provide additional benefits as wheel access was limited to one mouse at a time.

In addition, variability in the activity of specific neural networks has been linked to mouse wheel activity (Rhodes et al., 2003): lateral hypothalamic orexin/hypocretin and GAD65 networks have emerged as drivers of locomotion initiation (Kosse et al., 2017; Karnani et al., 2020), suggesting that interindividual differences in network activity might explain variability in wheel running. In addition, other factors which were not captured in our experiment, might also have contributed to variability in OD-plasticity. This includes exploratory behaviours not captured by our setup, a higher intrinsic plastic ability of the brain enabling OD-plasticity more independently of external factors, which might be related to overall higher fitness levels as also suggested by younger mice running more than older mice (Bartling et al., 2017).

Together, our newly developed gRW-setup allowing to correlate individual mouse running behaviour with individual measures of experience-dependent V1-plasticity has demonstrated a striking correlation between mouse running activity and OD-plasticity, highlighting the importance of individual behavioural tracking for explaining experimental data. Thus, our observations add to the growing body of evidence that individual behavioural choices can strongly affect individual brain plasticity and thus should be analysed more carefully in future studies.

## Supporting information

Video 1 Mouse entry to gRW

Video 2 Mouse exit from gRW

Model 1

## Author contributions

Study design by C.S. Data collection and initial processing by C.S., J.S and M.G. Formal analysis by C.S. Data interpretation by C.S. and S.L. Original manuscript draft by C.S. Reviewing and editing of manuscript draft C.S. and S.L.

## Data Availability Statement

The data supporting the findings of this research are available on request to the corresponding author, pending a formal data-sharing agreement and approval from the local ethics committee.

## Acknowledgements

We thank DFG (SFB889, Project B05 to S.L.) and the Dorothea Schlözer-Program for funding. We thank Tobias Mühmer and Jan Hoffmann, for constructing the gated running wheel cage, and Christina Ahlbrecht for technical assistance and animal care.

---

**Video 1**. Video footage of a mouse entering the gated running wheel compartment via flipping a seesaw.

**Video 2**. Video footage of a mouse leaving the gated running wheel compartment via flipping a seesaw. Note that its cage mate is blocked from entering the compartment. **Model 1**. 3D model of gated running wheel components

**Figure S1**. Electronic circuit of gated running wheel setup

**Figure S2**. Ilustration of PCA analysis of wheel running parameters averaged across 7 days (A-C) and from individual days (D-F). A,D: Data transformed into coordinate system of principal component (PC) 1 and PC 2 and graphical representation of loadings. B,E: Loadings of all 5 PC C,F: Variability explained by each PC.

**Figure S3**. Comparison of wheel running parameters between no MD and MD mice. No significant difference was observed.

