## Supplementary figures and images for "More running causes more ocular dominance plasticity in mouse primary visual cortex: new gated running wheel setup allows to quantify individual running behaviour of group-housed mice"

### Figure S1

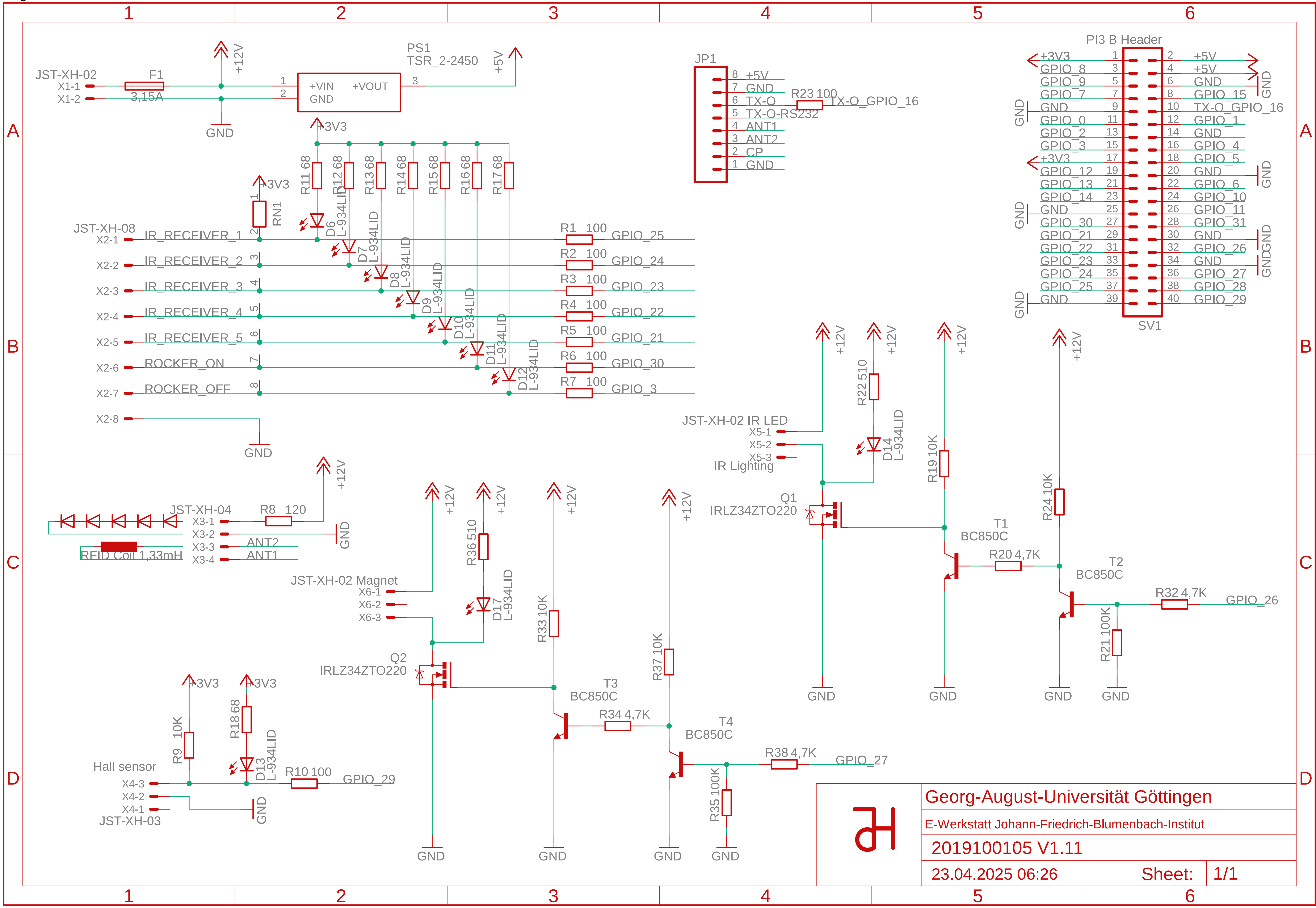

### Figure S2

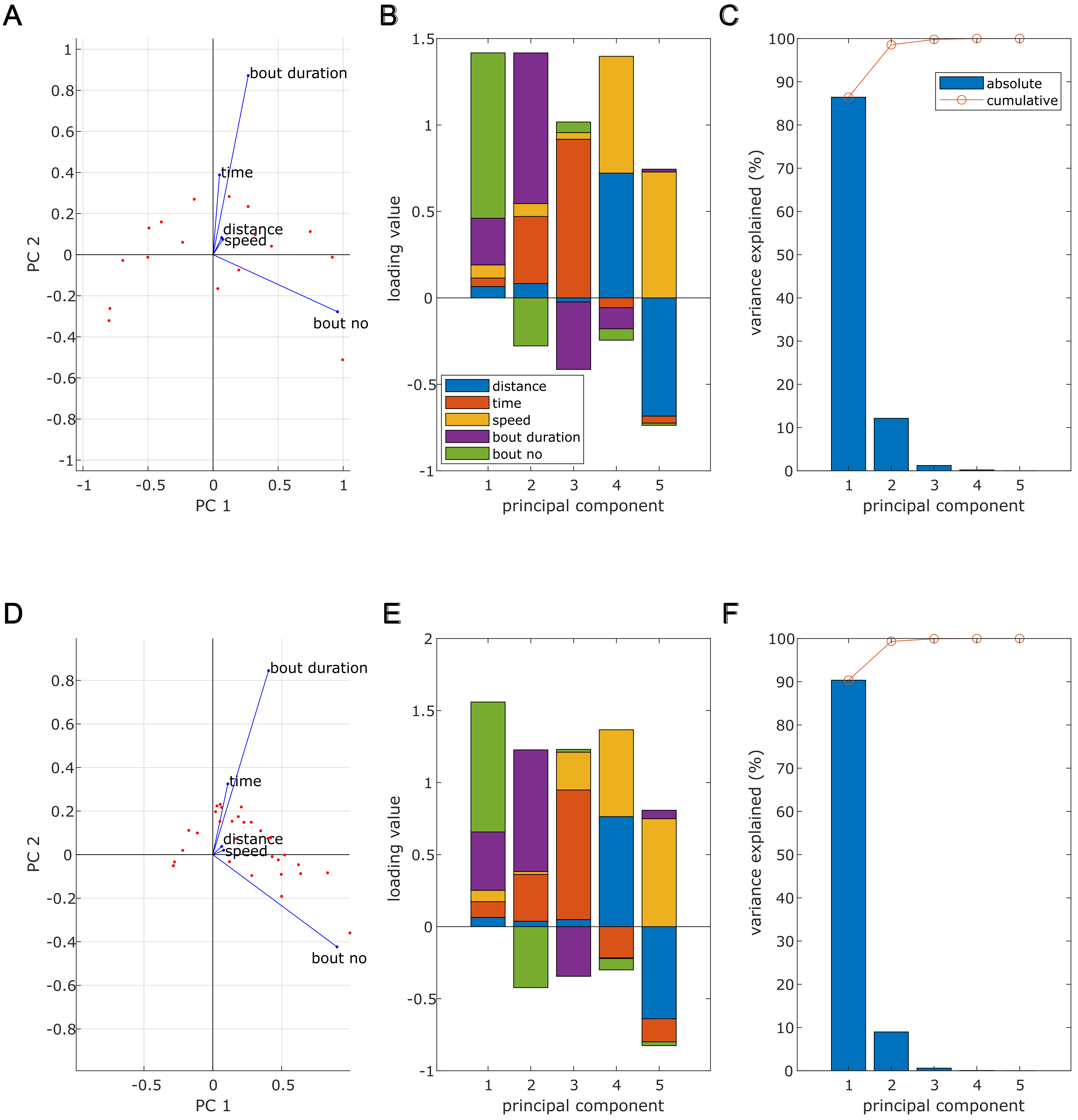

### Figure S3

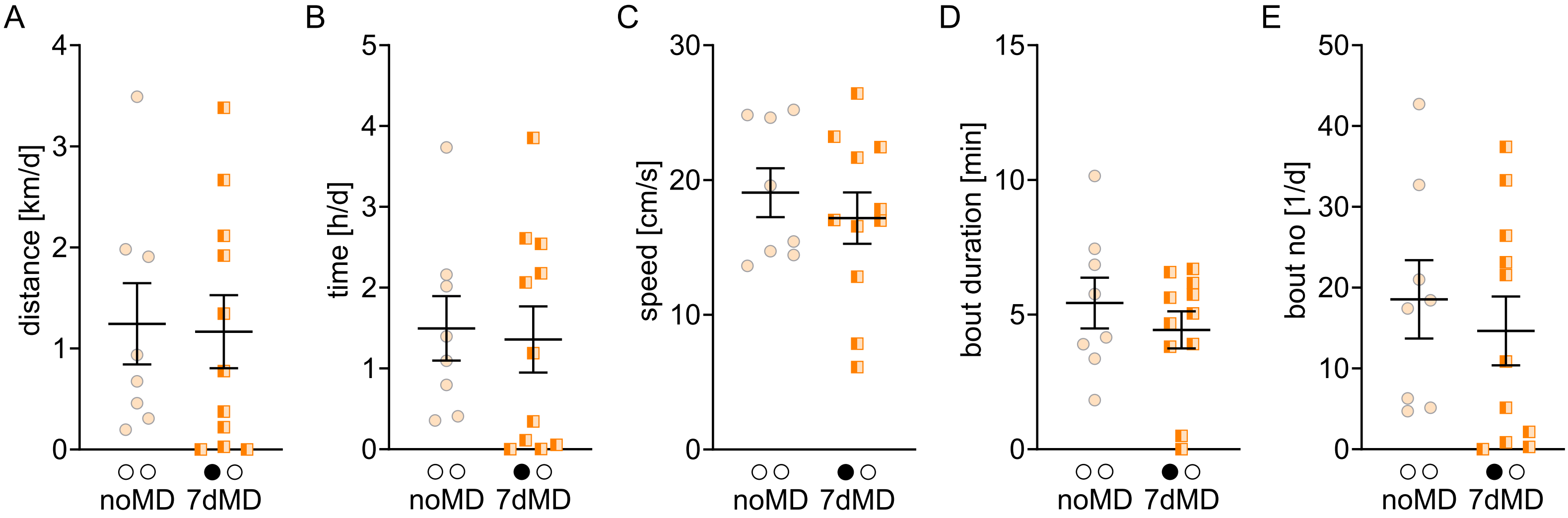
