## Supplementary material for "More running causes more ocular dominance plasticity in mouse primary visual cortex: new gated running wheel setup allows to quantify individual running behaviour of group-housed mice": Model 1

Figure 1-3: 3D-Model of gated running wheel components

| | |
|---|---|
| TITEL | |
| BAUTEILNR. | Auffangwanne |
| REVISION | |
| KONSTRUKTEUR | T. Mühmer |
| INGENIEUR | |

ANMERKUNGEN

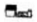

Primr

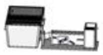

Standard

DIE INFORMATIONEN UND/ODER DAS MATERIAL IN DIESEM DOKUMENT SIND EIGENTUM VON BZW. VERTRAULICHE INFORMATIONEN UND/ODER VERTRAULICHES MATERIAL DES AUTORS. DIESE INFORMATIONEN DÜRFEN NICHT OHNE SCHRIFTLICHE GENEHMIGUNG VERWENDET, VERVIELFÄLTIGT, VERÖFFENTLICHT ODER OFFENGELEGT WERDEN. SIE SIND NUR FÜR IN DIESEM DOKUMENT ANGEGEBENE FERTIGUNGSOBJEKTE ZU VERWENDEN.
